# Overcoming cryo-EM map anisotropy reveals ALK-cytokine assemblies with distinct stoichiometries

**DOI:** 10.1101/2024.08.08.607122

**Authors:** Jan Felix, Steven De Munck, J. Fernando Bazan, Savvas N. Savvides

**Affiliations:** VIB-UGent Center for Inflammation Research, 9052 Ghent, Belgium; Unit for Structural Biology, Department of Biochemistry and Microbiology, Ghent University, 9052 Ghent, Belgium; ħ bioconsulting llc, Stillwater, MN, USA

## Abstract

Activation of Anaplastic lymphoma kinase (ALK) and leukocyte tyrosine kinase (LTK) by their cognate cytokines ALKAL2 and ALKAL1 play important roles in development, metabolism, and cancer. Recent structural studies revealed ALK/LTK-cytokine assemblies with distinct stoichiometries. Structures of ALK-ALKAL2 and LTK-ALKAL1 complexes with 2:1 stoichiometry determined by X-ray crystallography (De Munck *et al.* Nature, 2021) contrasted the 2:2 ALK-ALKAL2 complexes determined by cryo-EM (Reshetnyak *et al.,* Nature, 2021) and X-ray crystallography (Li *et al*., Nature, 2021). Here, we show based on reanalysis of the cryo-EM data deposited in EMPIAR- 10930 that over half of the ALK-ALKAL2 particles in the dataset are classified into 2D classes obeying a 2:1 stoichiometry besides the originally reported structure displaying 2:2 stoichiometry. Unlike particles representing the 2:2 ALK-ALKAL2 complex, particles for the 2:1 ALK-ALKAL2 complex suffer severely from preferred orientations that resulted in cryo-EM maps displaying strong anisotropy. Here, we show that extensive particle orientation rebalancing in cryoSPARC followed by 3D refinement with Blush regularization in RELION constitutes an effective strategy for avoiding map artefacts relating to preferred particle orientations and report a 3D reconstruction of the 2:1 ALK-ALKAL2 complex to 3.2 Å resolution from EMPIAR-10930. This new cryo-EM structure together with the crystal structures of ALK-ALKAL2 and LTK-ALKAL1 complexes with 2:1 stoichiometry reconciles a common receptor dimerization mode for ALK and LTK and provides direct evidence for the presence of an ALK-ALKAL2 complex with 2:1 stoichiometry next to the reported 2:2 stoichiometric assembly in the EMPIAR-10930 dataset. Finally, our analysis emphasizes the importance of public deposition of raw cryo-EM data to allow reanalysis and interpretation.

## Introduction & Results

Anaplastic lymphoma kinase (ALK) and the related leukocyte tyrosine kinase (LTK) are receptor tyrosine kinases (RTKs) that are activated by binding their cognate cytokines, ALKAL1 and ALKAL2 (also called FAM150A/B or AUGβ/α)[1–3]. Their signaling outputs elicit a range of pleiotropic activities in development and metabolism, and their dysregulated intracellular kinases are keenly pursued as therapeutic drug targets in various cancers[4]. However, the field had long been hampered by a paucity of insights into the intricate folds of ALK and LTK and their cytokine-bound complexes. A triad of recent studies[5–7] have aimed to provide the missing structural and mechanistic framework for the cytokine-mediated activation of ALK family receptors but diverged in their proposed assemblies.

Our study by De Munck *et al.*[5] provided representative assemblies for the entire ALK family by reporting crystal structures of both the ligand-binding extracellular regions of human ALK and LTK (ALKTG/LTKTG) in complex with ALKAL2 and ALKAL1, respectively (PDB entries 7nwz and 7nx0). These structures revealed how a single cytokine molecule nucleates homodimeric receptor assembly via distinct site 1 and 2 interaction interfaces, creating a key site 3 receptor-receptor contact (Supplementary Figure 1A). In contrast, Li *et al.*[6] and Reshetnyak *et al.*[7] have focused only on ALK-ALKAL2 complexes comprising the extracellular ligand-binding fragment of ALK and the membrane-proximal EGFL domains (ALKTG-EGFL), and reported assemblies obeying a 2:2 stoichiometry obtained via X-ray crystallography using an ALKTG-EGFL-ALKAL2 fusion construct (PDB entry 7LS0) or separate ALKTG-EGFL and full-length ALKAL2 via electron cryo-microscopy (cryo-EM, PDB entry 7n00), respectively (Supplementary Figure 1B). In this side-by-side architecture, the main cytokine-receptor interface corresponds to the high affinity site 1 also reported by De Munck *et al*.[5]. An additional ALK-ALKAL2’ interaction, covering 250 Å^2^ of buried surface area was proposed to be responsible for bridging the two 1:1 ALK-ALKAL2 subcomplexes into an assembly that could support signaling. Intrigued by the 2:2 stoichiometry differing from our 2:1 stoichiometric crystal structures of truncated ALKTG in complex with ALKAL2 (and LTKTG with ALKAL1), we closely examined the cryo-EM data and analysis reported by Reshetnyak *et al*.[7] as deposited in the EMPIAR data base (entry EMPIAR-10930).

At the outset, we identified a number of 3D-classes (Reshetnyak *et al*., Supplementary Figure 4 in reference [7], classes 1, 3, 7, 8 and 9) resembling the compact 2:1 ALKTG/LTKTG-cytokine complexes we had reported[5]. To investigate this apparent particle heterogeneity, we leveraged the raw cryo-EM data and uncleaned particle stack as deposited by Reshetnyak *et al.* (access code <u>EMPIAR-</u> <u>10930</u>). An initial data processing strategy following the methods described in Reshetnyak *et al*., including 8x binning of the data (Materials and Methods) revealed that 6 of the 10 most populated classes after initial 2D classification do not correspond to classes shown in the final 2D classification reported in Reshetnyak *et al.* (Figure 1A). This prompted us to pursue extensive data processing (Materials and Methods, Supplementary Figure 2) of particles present in both sets of 2D classes, namely the ones that correspond to clear 2-fold symmetry (Figure 1A, orange squares) and the ones that lack apparent symmetry (Figure 1A, blue squares). However, the final refined map of the C1 symmetric particles displayed severe preferential particle orientations leading to a map that is smeared along the Z-direction, a common problem in cryo-EM requiring sample and data collection optimization. Although the resulting map clearly displays two ALKTG-EGFL extracellular domains and one ALKAL2 copy in a similar 2:1 assembly as observed in the crystal structure of the ALKTG-ALKAL2 complex (Supplementary Figure 2), artefacts resulting from severe anisotropy hampered confident model building and prohibited detailed structural interpretations.

**Figure 1:**
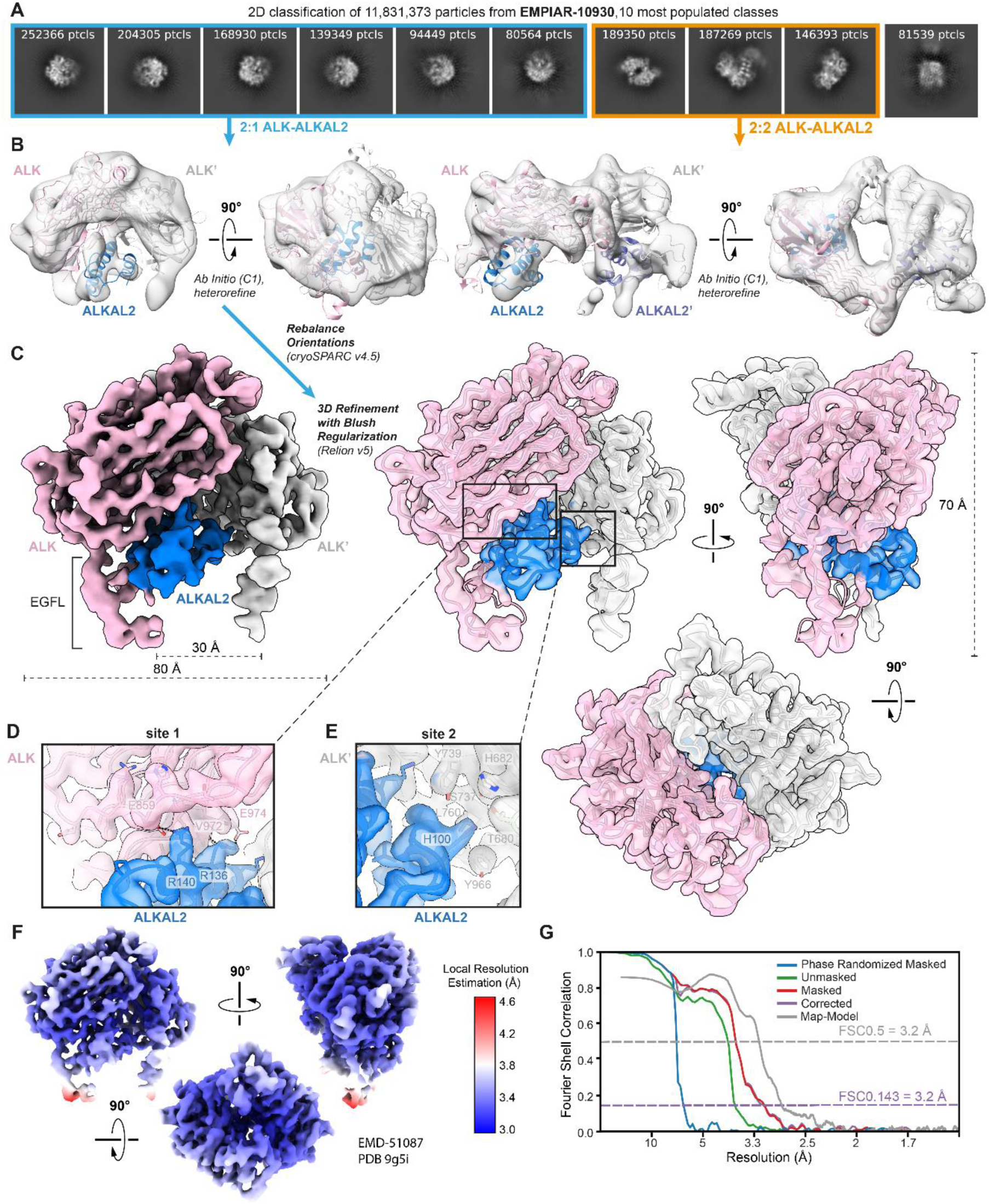
Reanalysis of cryo-EM data from EMPIAR entry EMPIAR-10930 deposited by Reshetnyak *et al.,* Nature (2021). (A) 10 most populated 2D class averages after performing 2D classification on 11.8 million selected particles. Classes corresponding to 2:1 ALK–ALKAL2 particles are highlighted in blue, while classes corresponding to 2:2 ALK–ALKAL2 particles are highlighted in orange. (B) *Ab initio* model generation in cryoSPARC[8] results in classes corresponding to both a 2:1 ALK-–ALKAL2 and 2:2 ALK–ALKAL2 assembly. Hetero-refinement of the 2:1 and 2:2 ALK–ALKAL2 classes results in the two models displaying 2:1 and 2:2 ALK–ALKAL2 assemblies, respectively. The ALK–ALKAL2 complexes with 2:1 (PDB 7nwz) and 2:2 (PDB 7n00) stoichiometries are shown fitted inside the maps, with ALK colored pink/grey and ALKAL2 colored blue/purple. (C) 3D refinement with Blush regularization[9] in RELION v5 of a trimmed particle set after extensive automatic rebalancing of particle orientations as implemented in cryoSPARC 4.5 (see also Materials and Methods & Supplementary Table 1). A front view is shown of a map sharpened in RELION using a B- factor of -100 Å^2^, with ALK colored pink/grey and ALKAL2 colored blue. Front, side and top views are shown of a transparent map, sharpened using deepEMhancer, with fitted structural model. (D & E) Zoomed-in insets illustrating the quality of map density corresponding to the two major interaction interfaces (site 1 & 2) in the ALK:ALKAL2 complex. Key interacting residues are annotated and shown as sticks. (F) Local Resolution Estimation performed in RELION v5 of the final map for the 2:1 ALK-ALKAL 2 complex. (G) Gold-standard Fourier Shell Correlation (FSC) plot corresponding to the final map of the 2:1 ALK:ALKAL2 complex. The corrected FSC (purple lines) is calculated after performing correction by noise substitution[31], and the resolution at FSC = 0.143 is annotated via a dotted purple line. A map-to model FSC curve calculated using the fitted model and an unsharpened, unfiltered full map is shown (grey line) with the resolution at FSC=0.5 annotated via a dotted grey line.

To overcome the apparent anisotropy presented in the 3D reconstructions towards a 2:1 ALKTG- EGFL–ALKAL2 complex, we reasoned that combining exhaustive automatic particle orientation rebalancing (available in cryoSPARC[8], v4.5) to exclude particles from over-populated direction bins, followed by 3D refinement with Blush regularization[9] in RELION v5, might result in more isotropic 3D reconstructions. Blush regularization uses a pre-trained denoising convolutional neural network on pairs of half-set reconstructions as a smoothness prior and is shown to remove artifactual densities resulting from uneven angular distributions or streaky features in overfitted maps[9]. This new regularization approach to resolving vexing cryo-EM structural data was successfully employed to reveal the distinct binding modes of the anti-CRISPR repressor Aca2 with DNA and RNA[10].

Starting from a set of selected 3D classes (1.4 million particles) after 3D hetero-refinement (Materials and Methods), we divided these 3D classes in two sets based on the absence or presence of apparent twofold symmetry. Such grouping resulted in 3D classes resembling either the C1- symmetric 2:1 ALKTG-ALKAL2 crystal structure reported by De Munck *et al*.[5] (876.806 particles, ~60% of the total), or the deposited C2-symmetric 2:2 ALKTG-EGFL-ALKAL2 cryo-EM map reported by Reshetnyak *et al.*[7] (523.467 particles, ~40% of the total) (Figure 2A).

**Figure 2:**
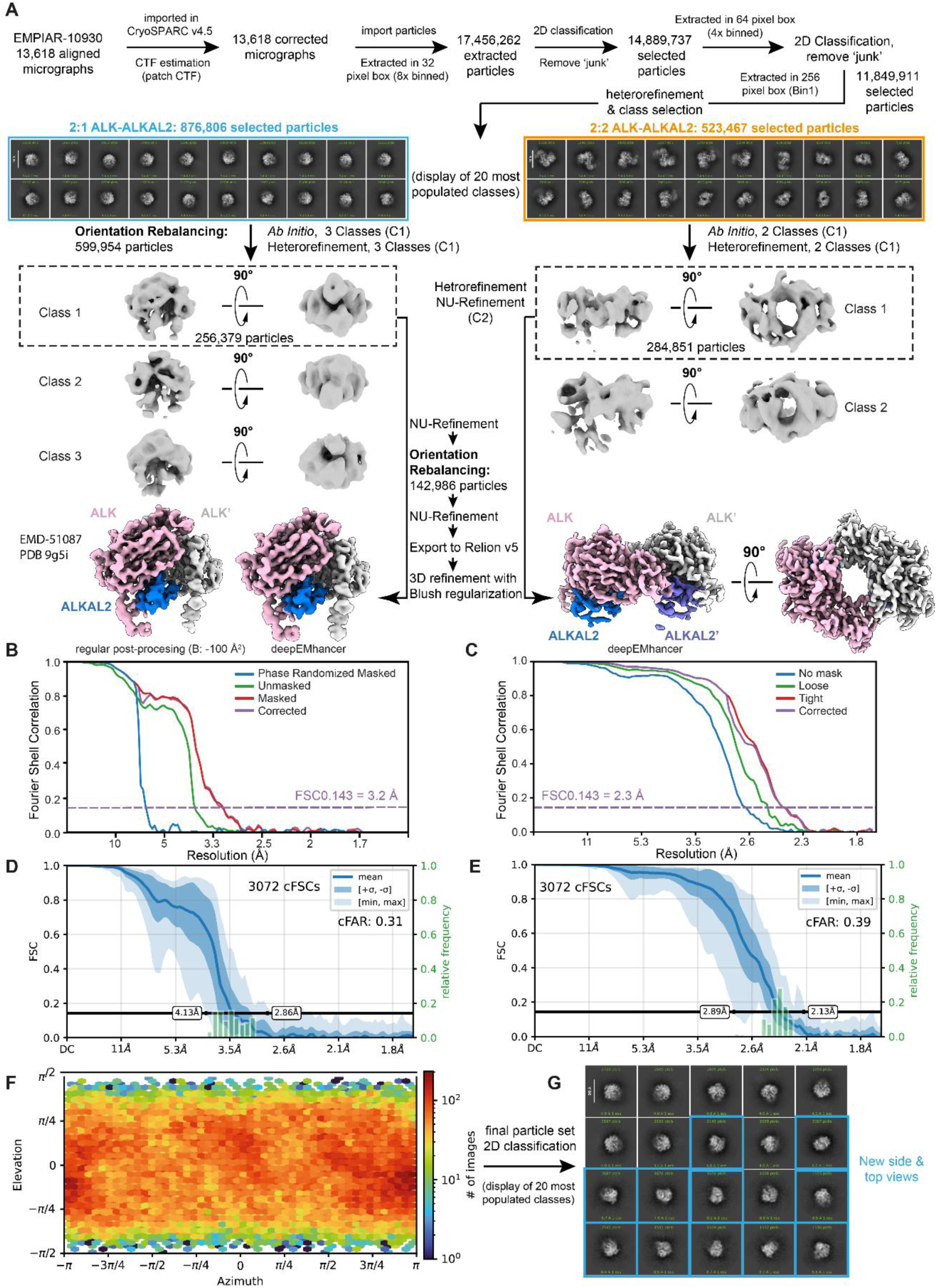
Final processing pipeline of cryo-EM dataset EMPIAR-10930. (A) Final processing steps were performed in CryoSPARC[8] v4.5 or RELION[22, 23] v5. Post-processing was performed in RELION or using deepEMhancer[25]. (B & C) Gold-standard Fourier Shell Correlation (FSC) plots corresponding to the final maps of the 2:1 (B) and 2:2 (C) ALK:ALKAL2 complexes. The corrected FSCs (purple lines) are calculated after performing correction by noise substitution[31], and the resolution at FSC = 0.143 is annotated via dotted purple lines. (D & E) conical FSC summary plots corresponding to the final maps of the 2:1 (D) and 2:2 (E) ALK:ALKAL2 complexes, generated via ‘Orientation Diagnostics’ in cryoSPARC v4.5. (F) Azimuth plot showing the distribution of orientations over Azimuth (X-axis) and Elevation (Y-axis) angles for the particle set corresponding to the final map of the 2:1 ALK-ALKAL2 complex, obtained after Orientation Rebalancing in cryoSPARC v4.5 and 3D refinement with Blush regularization[23] in RELION v5. (G) Display of the 20 most populated classes after 2D classification of the final particle stack corresponding to the 2:1 ALK-ALKAL2 complex. New side & top views not observed in initial 2D classification runs are shown as blue squares.

Firstly, we wondered whether our approach would lead to a cryo-EM map of similar resolution and informational content for the C2 symmetric ALKTG-EGFL-ALKAL2 complex with 2:2 stoichiometry as reported by Reshetnyak *et al.*[7]. Indeed, further data processing in cryoSPARC resulted in a 2.3 Å resolution map for the 2:2 ALKTG-EGFL-ALKAL2 complex (Materials and Methods). The resulting map has a comparable resolution as the deposited map by Reshetnyak *et al. (*EMD-24095, 2.3 Å) and, importantly, does not contain any particles from 2D classes corresponding to a 2:1 ALKTG-EGFL- ALKAL2 stoichiometry (Figure 2A). We note that, similarly as the cryoSPARC refined map reported in Reshetnyak *et* al., map density for the EGFL domains is less defined than for the RELION refined map obtained by Reshetnyak *et al.* at slightly lower resolution.

We then pursued analysis of the particle set with apparent resemblance to ALKTG-EGFL-ALKAL2 complexes with 2:1 stoichiometry (Materials and Methods). The combined use of Orientation Rebalancing in CryoSPARC and 3D refinement in RELION v5 with enabled Blush regularization, resulted in a massively improved final map at 3.2 Å resolution (Figure 1, Figure 2A, Supplementary Table 1) displaying an impressive improvement in isotropy, as evidenced by the 10-fold increase in the cFAR score (Figure 2D). Most importantly, post-processing in RELION yielded a sharpened map with clear main chain connectivity and density for most side chains, allowing complete model building of a 2:1 ALKTG-EGFL-ALKAL2 complex that includes the membrane-proximal EGFL-like domains of ALK. While the majority of the 2:1 ALKTG-EGFL-ALKAL2 cryo-EM map has a local resolution to better than 3.5 Å resolution (Figure 1F), the C-terminal tips of the EGFL domains are only resolved at 4 – 4.5 Å resolution pointing to their inherent flexibility. We next wondered whether 3D refinement with Blush regularization would be sufficient to obtain an isotropic map from the particle set without Orientation Rebalancing. Starting from the same particle set before the first Orientation Rebalancing step, followed by *Ab-Initio* model generation (3 classes), hetero-refinement and a final 3D refinement with Blush Regularization of the highest resolution class, resulted in a map at higher resolution (2.7 Å) but with lower isotropy (cFAR score of 0.1) than the map obtained by combining Orientation Rebalancing and 3D refinement with Blush regularization (Figure 3). We note that 3D refinement with Blush regularization of the non-Orientation Rebalanced particle set (cFAR score of 0.1) is superior in improving map isotropy than NU-refinement without or with Orientation Rebalancing, which yields maps with cFAR scores of 0.01 and 0.8 respectively (Figure 3). Thus, our analysis demonstrates that the combination of extensive Orientation Rebalancing and 3D refinement with Blush regularization may serve as a suitable strategy to overcome severe preferred particle orientations *In Silico*, albeit at the cost of some moderate loss in resolution.

**Figure 3:**
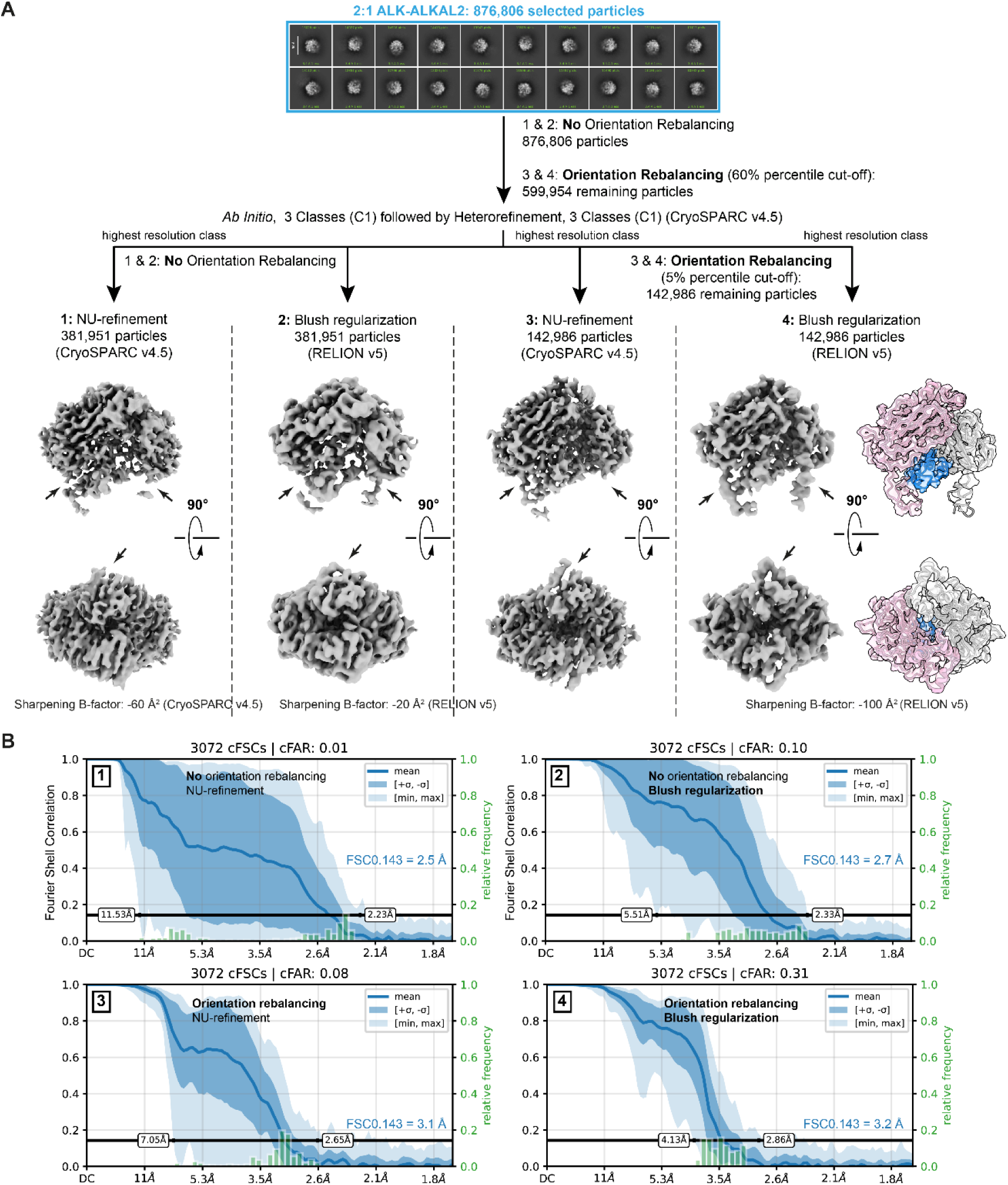
Comparison of map (an)isotropy following data processing strategies with or without Orientation Rebalancing and/or 3D refinement with enabled Blush Regularization. (A) Processing workflows (1 to 4) of cryo-EM dataset EMPIAR-10930 starting from 876.806 selected particles as outlined in Methods. Workflow 1 does not include Orientation Rebalancing (CryoSPARC v4.5) nor 3D refinement with Blush regularization (RELION v5). Workflow 2 is the same as 1 but with a final 3D refinement with Blush regularization in RELION v5 instead of NU-refinement in CryoSPARC v4.5. Workflow 3 utilizes Orientation Rebalancing in CryoSPARC v4.5 followed by NU-refinement in CryoSPARC, while workflow 4 combines Orientation Rebalancing in CryoSPARC with 3D refinement with Blush regularization in RELION. (B) Conical FSC summary plots for the final 3D refinements obtained in processing workflows 1, 2, 3 and 4, generated via ‘Orientation Diagnostics’ in cryoSPARC v4.5.

## Discussion

The new structure of ALKTG-EGFL-ALKAL2 with 2:1 stoichiometry presented here is a critical contribution to the present collection of ALK/LTK-cytokine complexes revealed by X-ray crystallography and cryo-EM. The elucidation of an ALKTG-EGFL-ALKAL2 complex with 2:1 stoichiometry based on a dataset that was initially reported to only support ALKTG-EGFL-ALKAL2 complexes with 2:2 stoichiometry underscores the power of modern cryo-EM to harness the conformational and stoichiometric diversity harbored in protein samples. Furthermore, our analysis emphasizes the importance of public deposition of raw (cryo-EM) data to allow reanalysis and interpretation. Most importantly, though, this new structure is now the most complete ALK- ALKAL2 complex with 2:1 stoichiometry to date and agrees closely with the ALKTG-ALKAL2 with 2:1 stoichiometry reported by De Munck et al.[5] featuring site 1 (ALK-ALKAL2), site 2 (ALK’-ALKAL2), and site 3 (ALK-ALK’) interaction interfaces (Figure 1). Accordingly, the root-mean-square deviation (r.m.s.d.) between these two structures is 1.3 Å over 675 aligned Cα-atoms. The site 2 interface present in the 2:1 ALK:ALKAL2 complex, but not in the 2:2 ALK-ALKAL2 complex, involves the α1 helix of ALKAL2 with a key histidine residue (H100) interacting with S737, Y739 and L760 of ALK’ (Figure 1E). Furthermore, the C-termini of the EGFL domains of the two ALK copies in the 2:1 ALKTG- EGFL-ALKAL2 complex obtained from EMPIAR-10930 are spaced ~30 Å apart (Figure 1), a distance that would allow facile juxtapositioning of receptor chains and TM helices as already illustrated for many RTK—cytokine assemblies featuring cytokine-mediated receptor dimerization such as KIT– SCF[11, 12], CSF-1R–CSF-1[13], Flt3–Flt3R[14], VEGF-VEGFR[15], PDGF-PDGFR[16], EGF–EGFR[17], HER2–HER3–NRG1β[18] and Insulin bound to insulin receptor (IR) family receptors[19, 20]. The presence of two distinct stoichiometries of the ALK-ALKAL2 complex among the particles imaged in the EMPIAR-10930 dataset is intriguing and their role in ALK signaling is currently unclear. One hypothesis could be that increasing local cytokine concentrations could shift an ALK-ALKAL2 assembly from a 2:1 to a 2:2 stoichiometry, resulting in an increase in the distance between the membrane proximal EGFL domains from 30 Å tot 90 Å, possibly leading to different signaling outcomes(Supplementary Figure 3). Interestingly, designed ligands for erythropoietin receptor aimed at tuning the distance between receptor monomers upon dimerization result in modulation of erythropoietin receptor signaling and phosphorylation efficiency of downstream adaptors[21]. Finally, structural, biophysical and cellular studies, including full-length ALK-ALKAL2 complexes, will be required to fully define the possible roles of distinct ALK/LTK-cytokine stoichiometries.

## Materials and Methods

### Reanalysis of cryo-EM data as reported in EMPIAR-10930

Initial data analysis was carried out in CryoSPARC[8] v3.3.1, and additional steps were performed in either cryoSPARC[8] v4.5 or RELION[22, 23] v5, specifically for the use of Orientation Rebalancing (‘Rebalance Orientations’ job) and 3D refinement including Blush regularization[9] respectively. Dose-weighted and motion-corrected micrographs were downloaded from https://www.ebi.ac.uk/empiar/EMPIAR-10930/ and were CTF-corrected using patch CTF estimation. After importing the uncleaned particle stack (~18 million particles) from EMPIAR-10930 and extracting the corresponding particles (8x binned), several iterative cycles of reference-free 2D classification were performed to get rid of ‘junk’ particles, resulting in a ‘cleaned’ set comprising ~11.8 million particles. This number of particles approaches the total number of particles (~12.4 million) used by Reshetnyak *et al*. Particle unbinning followed by reference-free 2D classification revealed that almost 2/3 of the ten most populated classes do not correspond to classes shown in the final 2D classification represented in Reshetnyak *et al.* (Figure 1a). *Ab initio* models generated on a selected subset of most populated classes (4x binned, 1.5 million particles, 3 *Ab Initio* classes, C1 symmetry) followed by a round of heterorefinement revealed one model that clearly corresponds to the 2:2 ALK–ALKAL2 structure reported by Reshetnyak *et al.,* as well as a model resembling the 2:1 ALK–ALKAL2 structure presented in De Munck *et al.* (Figure 1b & Supplementary Figure 2). Next, the three *Ab Initio* classes were used together with the cleaned dataset of ~11.8 million particles (unbinned), as input for heterorefinement (C1, 9 classes). Class 1 and 2 of this heterorefinement were grouped together, and selected particles (~1.2 million) were used for 3D classification without alignment (8 classes). Class 5 after 3D classification was selected (334,654 particles) and used as an input for a final Non-Uniform (NU) refinement[24], While the final map shows the presence of two ALK copies and one ALKAL2 copy (with its characteristic three central α-helices), the map suffers from the severe preferential particle orientations of the 2:1 ALK–ALKAL2 particles, as evidenced by the paucity of different views in the 2D classes as well as the Azimuth versus Elevation plot shown in Supplementary Figure 2. Indeed, Orientation Diagnostics, as implemented in the newest version of cryoSPARC[8] (v4.5), pointed to extreme anisotropy corresponding to a cFAR score of 0.03 (Supplementary Figure 2). Accordingly, the anisotropy present in the obtained NU-refined map of the 2:1 ALKTG-EGFL–ALKAL2 complex prohibited further model building and refinement.

Next, starting from the heterorefinement of 11.849.911 particles, highest resolution 3D classes were grouped in two pools based on the presence or absence of twofold symmetry. Further processing of the particles with twofold symmetry (523.467 particles) included one additional round of 2D classification to remove remaining junk classes, *ab-Initio* model generation (2 classes), hetero-refinement and 3D NU-refinement of the highest resolution class while applying C2 symmetry, and resulted in a 2.3 Å resolution map for the 2:2 ALKTG-EGFL-ALKAL2 complex (Figure 2). Further processing of the particles without apparent symmetry (876.806 particles) by 2D classification showed that 2D classes mainly consisted of side views of the 2:1 ALK-ALKAL2 complex, with a lack of discernible top views (Figure 2). We next used Orientation Rebalancing in cryoSPARC v4.5 to trim overpopulated views in the particle stack using a rebalance percentile threshold of 60, resulting in 599.954 remaining particles. These particles were used as an input for an Ab Initio refinement job (C1, 3 classes) followed by heterorefinement (C1, 3 classes). The highest resolution 3D class (class 1) was used for NU-refinement, followed by another round of Orientation Rebalancing, now using a more stringent rebalance percentile threshold of 5. Retained particles (142.986) where used for a second NU-refinement and subsequently exported to RELION v5 for a final round of 3D refinement with enabled Blush regularization. A final 2D classification was performed on the final particle stack (142.986 particles) to demonstrate the appearance of new side and top views not seen in previous 2D classification runs (Figure 2). Orientation Diagnostics jobs were run on obtained intermediate 3D reconstructions as well as the final map to monitor improvements in map isotropy. The final obtained map has a Gold-Standard Fourier Shell Correlation (FSC) resolution of 3.2 Å at the 0.143 threshold (Figure 1, Supplementary Table 1), and was post-processed either in RELION using a sharpening B-factor of -100 Å^2^, or by using deepEMhancer[25].

### Model Building & Refinement

Two copies of ALKTG-EGFL and one copy of ALKAL2 were extracted from PDB ID 7n00, and rigid-body fitted in the sharpened map of the 2:1 ALKTG-EGFL-ALKAL2 complex (B-factor: -100 Å^2^) using USCF Chimera[26]. The resulting rigid-body fitted model was subjected to a cycle of manual building in Coot[27], aided by both the regularly sharpened map and a map sharpened using DeepEMhancer[25]. Next, the manually-built and corrected model was real-space refined against the regularly sharpened map in Phenix v1.19.2-4158[28], using global minimization, local grid search, atomic displacement parameter (ADP) refinement, secondary structure restraints and Ramachandran restraints. Several cycles of manual building in Coot were followed by real-space refinement in Phenix. After the last cycle of manual building in Coot, a nonbonded-weight parameter of 200 was used during refinement in Phenix. The resulting model was subjected to a final refinement against the two half-maps in REFMAC-Servalcat[29] run within CCP-EM suite v1.6[30], with automatic weighting applied. The final model has a map-model FSC of 3.20 Å at the 0.5 threshold, calculated using the unfiltered, unsharpened full map (Figure 1). A summary of cryo-EM data collection, processing, refinement and validation statistics can be found in Supplementary Table 1.

## Data Availability

The cryo-EM map and accompanying structural model for the 2:1 ALKTG-EGFL-ALKAL2 complex have been deposited in the EMDB/PDB with accession codes: EMD-51087/9g5i. The particle .star file obtained after 3D refinement with Blush Regularization in RELION 5 is added to this manuscript as a Supplementary Data file, and can be used to extract particles after running CTF estimation on the Motion Corrected and Dose Weighted (*_DW.mrc) micrographs available from EMPIAR-10930 (https://www.ebi.ac.uk/empiar/EMPIAR-10930/).

## Supporting information

Particle .star file obtained after 3D refinement with Blush Regularization in RELION 5

## Acknowledgments

S.N.S. acknowledges research support from the FWO (grant number G0B4918N) and the Flanders Institute for Biotechnology (VIB, grant number C0101).

## Author contributions

J.F and S.N.S. wrote the manuscript with contributions from all authors. J.F performed analysis and processing of the deposited EMPIAR-10930 cryo-EM dataset with contributions from S.D.M. J.F. and S.D.M. performed structural analyses with contributions from S.N.S. and J.F.B.

## Supplementary Information

**Supplementary Figure 1:**
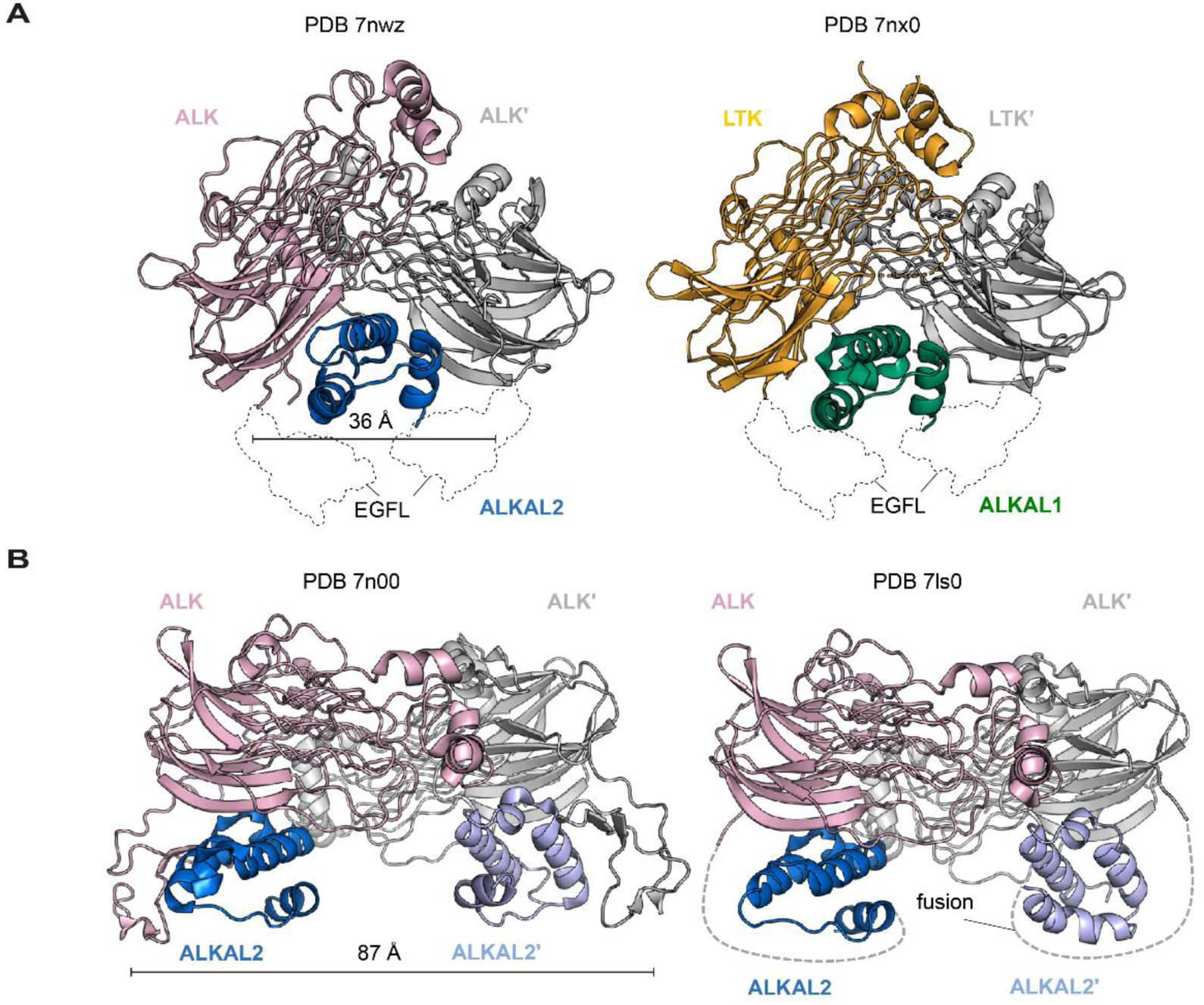
Comparison of 2:1 ALK/LTK and 2:2 ALK receptor-ligand complexes. (a) Cartoon views of structures of ALK-ALKAL2 and LTK-ALKAL1 complexes with 2:1 stoichiometry as determined by De Munck *et al*.[5] (PDB entries 7nwz and 7nx0) and (b) of ALK-ALKAL2 with 2:2 stoichiometry as determined by Reshetnyak *et al.*[7] and Li *et al.*[6] (PDB entries 7n00 and 7ls0). Putative positions of the EGFL domains absent in (a) are shown as dotted outlines.

**Supplementary Figure 2:**
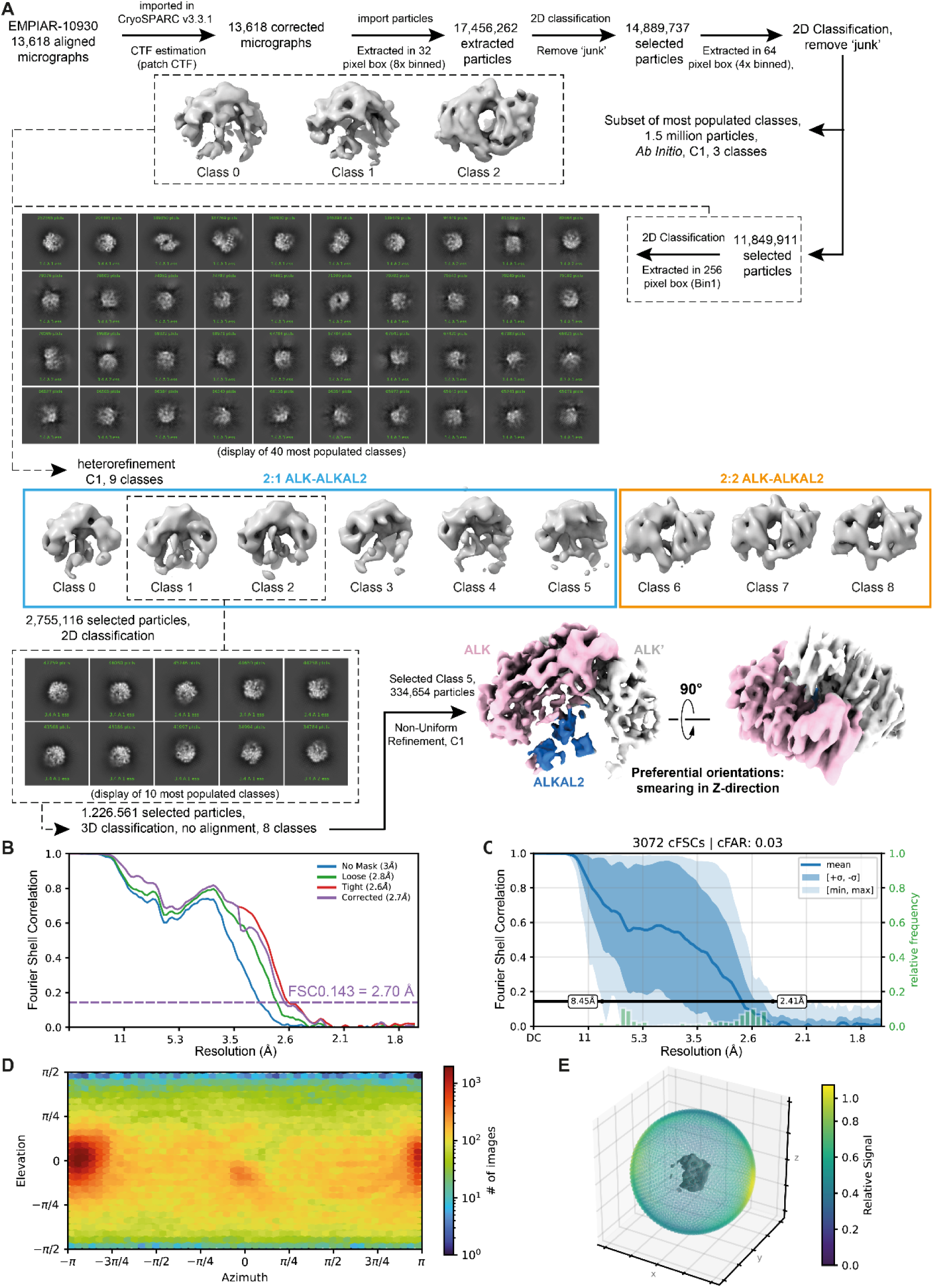
Initial processing workflow of cryo-EM dataset EMPIAR-10930. (A) All initial processing steps were performed in CryoSPARC[8] v3.3.1. (B) Gold-standard Fourier Shell Correlation (FSC) plot corresponding to the final map obtained after non-uniform refinement (C1 symmetry). Curves are shown after applying no mask (blue), a loose mask (green), or a tight mask (red) to the two half maps before calculating FSCs. The corrected FSC (purple) is calculated using the tight mask with correction by noise substitution[31], and the resolution at FSC= 0.143 is annotated via a dotted purple line. (C) conical FSC summary plot generated via ‘Orientation Diagnostics’ in cryoSPARC v4.5. (D) Left: Azimuth plot showing the distribution of orientations over Azimuth (X-axis) and Elevation (Y-axis) angles for the particle set corresponding to the NU-refined map shown in (A). Right: Plot showing Relative Signal amount vs. Viewing Direction.

**Supplementary Figure 3:**
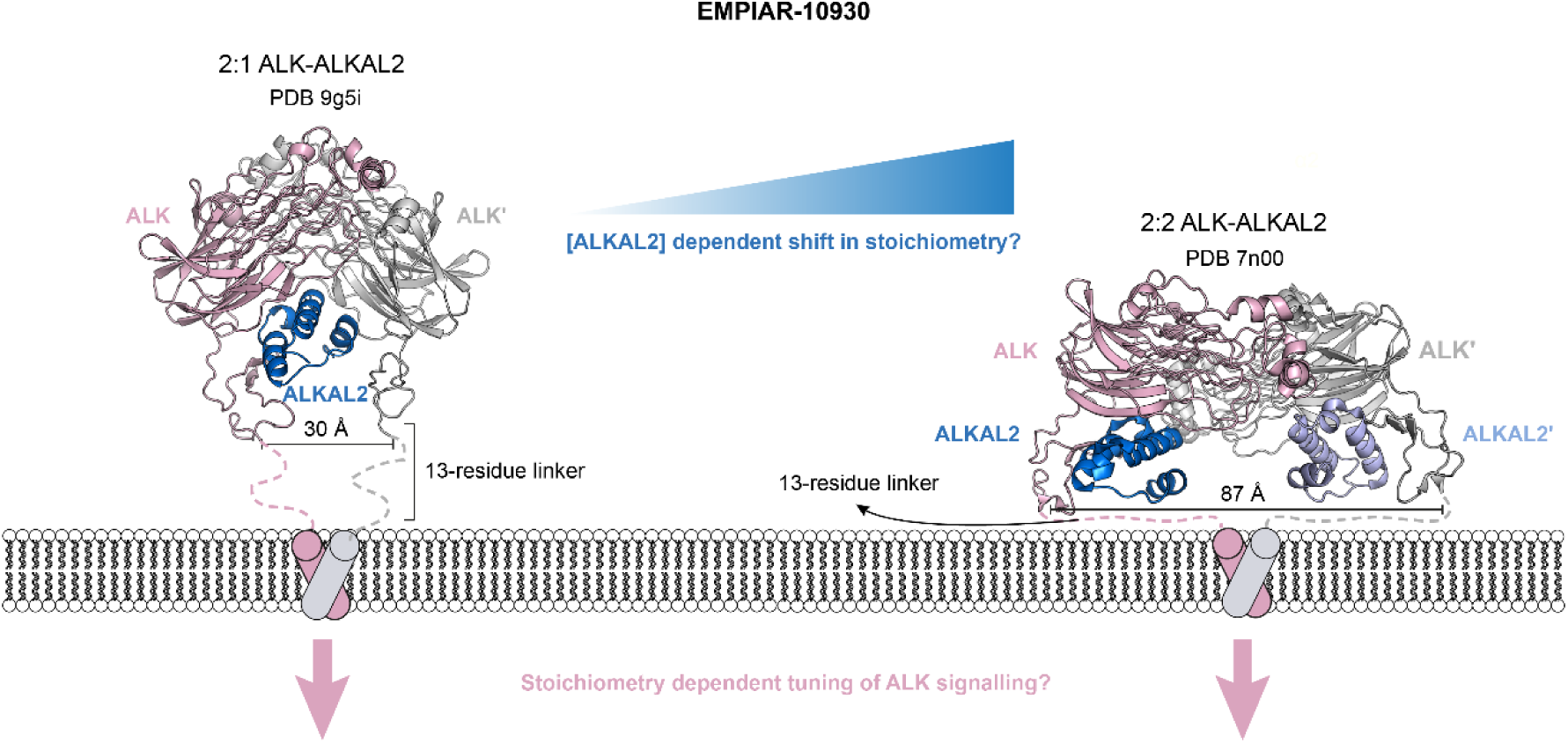
Side-by-side comparison of 2:1 and 2:2 ALK-ALKAL2 complexes present in the EMPIAR-10930 cryo-EM dataset. Models with pdb code 9g5i (this study) and 7n00 (Reshetnyak *et al*., 2021) are shown as cartoons with ALK/ALK’ colored pink/grey and ALKAL2 colored blue in the 2:1 ALK-ALKAL2 complex, or blue/purple in the 2:2 ALK-ALKAL2 complex. The distance between the C-termini of membrane- proximal EGFL domains is annotated.

**Supplementary Table 1.** Cryo-EM data collection, processing, refinement and validation statistics.

| 2:1 ALK<sub>TG</sub>-EGFL-ALKAL2 |  |
| --- | --- |
| **Data Collection (EMPIAR-10930, Reshetnyak *et al.* 2021[7])** |  |
| Magnification | 15,000 |
| Voltage (kV) | 300 |
| Electron exposure (e<sup>-</sup>/Å<sup>2</sup>) | 88.4 |
| Defocus range (μm) | 0.8 – 1.8 |
| Pixel size (Å) | 0.826 |
| Micrographs | 13.618 |
| **Data Processing (This study)** |  |
| Symmetry imposed | C1 |
| Initial particle images (no.) | 18,053,705 |
| Final particle images (no.) | 142,986 |
| Map resolution (Å) | 3.2 |
| FSC threshold | 0.143 |
| Map resolution range (Å) | 3.0 – 5.5 |
| **Refinement (This study)** |  |
| Initial model used (PDB code) | 7N00 |
| Model resolution (Å) | 3.2 |
| FSC threshold | 0.5 |
| Map sharpening B factor (Å<sup>2</sup>) | -100 |
| Model composition |  |
| Non-hydrogen atoms | 5,252 |
| Protein residues | 732 |
| B factors (Å<sup>2</sup>) |  |
| Protein | 165.45 |
| R.m.s. deviations |  |
| Bond lengths (Å) | 0.009 |
| Bond angles (°) | 1.193 |
| Validation |  |
| MolProbity score | 1.86 |
| Clashscore | 9.85 |
| Poor rotamers (%) | 0.00 |
| CaBLAM outliers (%) | 2.64 |
| CC (mask/volume) sharpened, unsharpened | 0.76/0.71, 0.80/0.88 |
| EMRinger |  |
| (sharpened/unsharpened) | 2.10/1.51 |
| Ramachandran plot |  |
| Favored (%) | 95.04 |
| Allowed (%) | 4.96 |
| Disallowed (%) | 0.00 |
| Ramachandran Z-score |  |
| Whole | -1.92 |
| Helix | -1.17 |
| Sheet | -1.29 |
| Loop | -1.25 |

